# HAX1 drives assembly and activation of the mitochondrial intermembrane space chaperone CLPB

**DOI:** 10.64898/2026.05.24.727526

**Authors:** Monifa A. V. Fahie, Julia G. Hoffman, Mimi S. Kay, Julia R. Kardon

## Abstract

The multicellular metazoan lineage acquired a novel chaperone in the mitochondrial intermembrane space, the AAA+ disaggregase and refoldase CLPB. Although it is not known how they function together, CLPB and the intrinsically disordered IMS protein HAX1 interact and share disease and cellular phenotypes; loss of function in either gene causes severe congenital neutropenia as well as neuropathology and causes many proteins in the IMS and its bounding membranes to become insoluble. Through biochemical reconstitution, we here find that HAX1 is a direct stimulatory cofactor of CLPB. HAX1 promotes oligomerization of CLPB into an active disaggregase and stimulates the ATPase and refoldase activities of the oligomeric complex. A short peptide within HAX1 is necessary for direct interaction with the ankyrin domain of CLPB, but stimulation of CLPB activity requires additional elements of HAX1. Characterization of CLPB and CLPB-HAX1 oligomers indicates that HAX1 shifts the predominant oligomeric state of CLPB from a dodecamer to an apparent hexamer elaborated with HAX1, suggesting that this smaller oligomer is important during the cycle of CLPB function with clients.

## INTRODUCTION

The mitochondrial intermembrane space (IMS) is a relay station for protein import to all mitochondrial compartments, in addition to coordinating electron transport complex assembly, mitochondrial dynamics, and signaling (Busch et al., 2023; Suomalainen and Nunnari, 2024). These processes generate a large load of proteins inhabiting partially folded states or assembly intermediates, leaving them vulnerable to misfolding and aggregation. At the origin of multicellularity in holozoans, the IMS acquired a novel protein chaperone, the AAA+ ATPase CLPB/Skd3(Erives and Fassler, 2015). Loss of CLPB function causes increased insolubility among many proteins in the IMS and its bounding inner and outer mitochondrial membranes (Baker et al., 2024; Cupo and Shorter, 2020; Fan et al., 2022), indicating that this multicellular-specific chaperone is important for maintaining IMS protein homeostasis. Human mutations in CLPB cause two rare and severe syndromes: severe congenital neutropenia (SCN), and a multisystem disorder 3-methylglutaconic aciduria type VII (MGCA7) which includes neuropathology and neutropenia (Saunders et al., 2015; Tucker et al., 2022; Warren et al., 2022; Wortmann et al., 2015), underscoring the importance of CLPB function.

CLPB is partly homologous to a bacterial disaggregase, ClpB, and related disaggregases in the cytoplasm (Hsp104) and mitochondrial matrix (Hsp78) of unicellular eukaryotes. These unicellular homologs share a AAA+ protein unfoldase domain with CLPB, but they additionally contain a second AAA+ domain as well as accessory domains. These domains are important for recruitment of aggregated substrates to ClpB and cooperation of ClpB with Hsp70/40 to resolubilize substrates (Glover and Lindquist, 1998; Goloubinoff et al., 1999; Krzewska et al., 2001). Metazoan CLPB was transformed by replacement of the N-terminal AAA+ domain and regulatory domains with an ankyrin-repeat domain (ANK), as well as by localization to the IMS (Erives and Fassler, 2015; Thevarajan et al., 2020). With these transformations, CLPB retained disaggregase activity, but acquired independence from Hsp70/40 and also an independent refoldase activity (Cupo and Shorter, 2020; Gupta et al., 2023). Like its unicellular homologs, CLPB forms a spiraling hexamer that engages substrates for unfolding at its central pore. The ANK domain projects above the substrate-recruiting side of the hexamer and can additionally make trans-hexamer contacts to form a dodecameric CLPB complex (Cupo et al., 2022; Gupta et al., 2023; Wu et al., 2023). Mutations that impair dodecamer formation impaired CLPB activity as a refoldase (Gupta et al., 2023), leading to the proposal that the fenestrated enclosure formed by the ANK domains in the dodecamer functions as a protective cage for substrate refolding. How the hexameric and dodecameric forms of CLPB contribute to function with its physiological partners in the IMS has not been tested.

The IMS-localized, intrinsically disordered protein HAX1 interacts with CLPB directly through the ANK domain(Fan et al., 2022; Zhang et al., 2025), is one of the most enriched binding partners of CLPB across interaction proteomic studies (Baker et al., 2024; Chen et al., 2019; Fan et al., 2022; Wakula et al., 2020; Wu et al., 2023), and becomes less soluble in cells lacking CLPB (Baker et al., 2024; Cupo and Shorter, 2020), suggesting that HAX1 is either a high-priority client of CLPB or a functional partner in its action on other IMS proteins. Supporting this latter idea, loss of HAX1 increases IMS and mitochondrial membrane protein insolubility similarly to loss of CLPB (Fan et al., 2022) and mutations in HAX1 cause a form of Kostmann syndrome, a severe congenital neutropenia with frequent neurologic symptoms (Klein et al., 2007), thus substantially overlapping in phenotype with disease alleles of CLPB. Here, we directly test how HAX1 affects CLPB activity in a purified system and find that it functions as an activating cofactor of CLPB.

## RESULTS

### HAX1 functions as a stimulatory cofactor with CLPB

To test the idea that HAX1 is a functional partner rather than a client of CLPB, we probed their behavior together *in vitro*. Previous studies of CLPB activity as a disaggregase and a refoldase have analyzed the longest splice form of CLPB (annotated as isoform 1) (Cupo et al., 2022; Cupo and Shorter, 2020; Gupta et al., 2023). Most cells, however, express the shorter isoform 2, which has a truncated linking element between second and third ankyrin motifs in the ANK domain. This isoform has been reported to bind HAX1 more tightly (Fan et al., 2022). Both isoforms contain the same presequence, which is processed by MPP and then PARL to generate mature CLPB (Saita et al., 2017). We therefore used the PARL-processed form of CLPB isoform 2 in our experiments, referred to as CLPB.

Because HAX1 is intrinsically disordered and has no known biochemical activity on its own, we asked whether it could affect the core activity of CLPB as a disaggregase. Addition of equimolar HAX1 stimulated CLPB to reactivate aggregated firefly luciferase (FFL_agg_), a model substrate for CLPB disaggregation (Fig. 1A, Fig. S1A). Because HAX1 was insoluble when recombinantly expressed, we tested both resolubilized HAX1 and HAX1 produced as a soluble fusion with a maltose binding protein (MBP) domain. Both preparations activated CLPB disaggregase (Fig. S1A), but MBP-HAX1 was more consistent in activity across preparations; we therefore used MBP-HAX1 in our later experiments (referred to as HAX1). The intrinsically disordered protein casein, which CLPB has previously been shown to engage as a client (Cupo et al., 2022; Gupta et al., 2023), did not stimulate CLPB disaggregase activity, nor did isolated MBP (Fig. 1A), indicating that the stimulatory effect of HAX1 was specific. We noted that in the absence of HAX1, CLPB disaggregase activity plateaus after 60 minutes, similar to previous observations with CLPB alone (Cupo et al., 2022; Gupta et al., 2023), whereas in the presence of HAX1, CLPB continues to reactivate FFL_agg_ over at least two hours. This difference in the duration of disaggregase activity was not due to general loss of ATPase activity; ATP hydrolysis without HAX1 is stable throughout the duration of the disaggregase assay (Fig. S1C). We also observed that HAX1 potently stimulated the ATPase rate of CLPB (Fig. 1B, Fig. S1B,C), compared to weak or absent stimulation by casein or FFL_agg_, respectively (Fig. 1B). Compared to model client proteins, the unique ability of HAX1 to stimulate CLPB as a disaggregase and its potent stimulation of ATP hydrolysis indicates that HAX1 specifically enhances CLPB enzymatic activity in a mechanism distinct from that of an unfolded or aggregated substrate.

**Figure 1.**
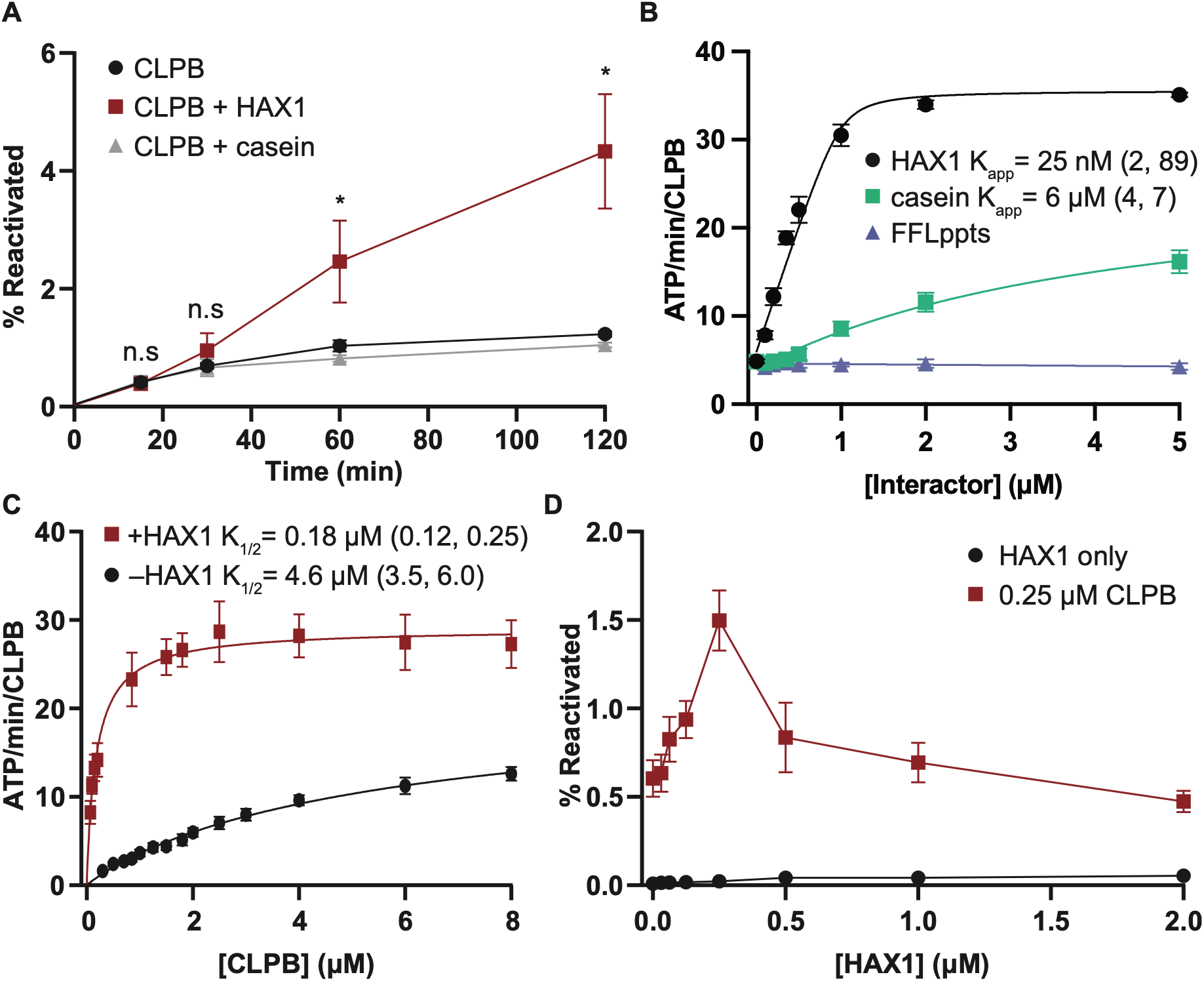
HAX1 stimulates CLPB disaggregase and ATPase activities by enhancing oligomerization. (A) Reactivation of firefly luciferase aggregates by 0.25 μM CLPB, ± 0.25 μM HAX1 or 0.25 μM model substrate FITC-casein, n = 2. (B) ATP hydrolysis rate of 1 μM CLPB in the presence of 0-5 μM HAX1, FITC-casein, or aggregated luciferase (FFLppts), fit to a quadratic velocity equation, n = 2. 95% confidence interval of fit in parentheses. Fit to FFLppts data was not statistically meaningful. The p value for K_app_ differing between HAX1 and casein <0.0001. (C) Oligomerization of CLPB, monitored using ATP hydrolysis as a proxy, fit to allosteric sigmoidal equation. HAX1, when present, is included at a 2:1 molar ratio with CLPB, n = 20 -HAX1, 10 +HAX1. p for K_1/2_ < 0.0001. (D) Reactivation of luciferase aggregates by 0.25 μM CLPB in the presence of 0-2 μM HAX1 (n = 6) or HAX1 alone (n = 3). For all plots, error = SEM; 95% confidence interval indicated for parameters extracted from fitted equations. Statistical significance denoted by p < 0.05 (*), p < 0.005 (**), p < 0.0005 (***).

We observed that HAX1 stimulation of disaggregation activity peaks at an equimolar HAX1:CLPB ratio and drops sharply as HAX1 concentration becomes superstoichiometric with CLPB (Fig. 1D). This inhibitory behavior suggests that once a specific binding site for HAX1 per ANK domain is saturated, additional HAX1 may compete with substrate for engagement by the CLPB pore. We also observe that HAX1 stimulation of CLPB disaggregase at equimolar concentrations becomes less significant and then inhibitory as CLPB concentration increases (Fig. S1D). A possible source of this effect may be that at higher total concentrations, ANK-domain bound HAX1 could compete with client in trans on another CLPB oligomer.

We additionally tested the effect of HAX1 on oligomerization of CLPB. Like other AAA+ ATPases, the ATPase active site of CLPB is formed at the interface of two protomers. CLPB is thus inactive as a monomer and gains ATPase activity on formation of hexamers and dodecamers, which appear to have similar ATPase activity (Gupta et al., 2023). We therefore monitored oligomerization of CLPB by proxy of its ATPase rate. HAX1 potently stimulated oligomerization of CLPB, decreasing the K_1/2_ for oligomerization by over 25-fold (Fig. 1C). Because HAX1 stimulation of CLPB disaggregase and ATPase activity was most apparent when CLPB concentration was well below its K_app_ (Fig. 1B-D), we attribute these effects primarily to the ability of HAX1 to promote oligomerization of CLPB. However, it is difficult to exclude the possibility that HAX1 stimulates the disaggregase activity of fully oligomerized CLPB as well, due to the apparent competition of HAX1 we observe with our model substrate at higher concentrations of the CLPB-HAX1 complex.

### A short helix of HAX1 is necessary for CLPB binding, but is insufficient to stimulate CLPB activities

HAX1 was previously demonstrated to bind to the ANK domain of CLPB in cells (Fan et al., 2022; Wu et al., 2023). To generate a more specific hypothesis for HAX1-CLPB contacts, we used AlphaFold3 to model a structure of their complex. This model contained a single high-confidence interaction (PAE < 5Å), positioning the predicted α3 helix of HAX1 (residues 125-130) in a groove in the CLPB ANK domain (Fig. 2A, B). To test the necessity of this predicted contact for activity, we truncated this region from HAX1 (HAX1^Δ125-130^). HAX1^Δ125-130^ did not stimulate disaggregase activity (Fig. 2C) and reduced the potency of ATPase stimulation to that observed for casein (Fig. 2D, Fig.1A). HAX1^Δ125-130^ did not bind CLPB (Fig. S2A), indicating that this predicted contact site is necessary for HAX1 to bind and activate CLPB. A mutation in this region, L130R, was found as part of a compound heterozygous genotype in a patient with severe congenital neutropenia (Lanciotti et al., 2010) and blocked interaction with CLPB in cells (Fan et al., 2022). This mutation also modestly decreased stimulation of the ATPase and disaggregase activity of CLPB by HAX1 and did not block interaction of the purified proteins (Fig. 2C,D; Fig. S2A).

**Figure 2.**
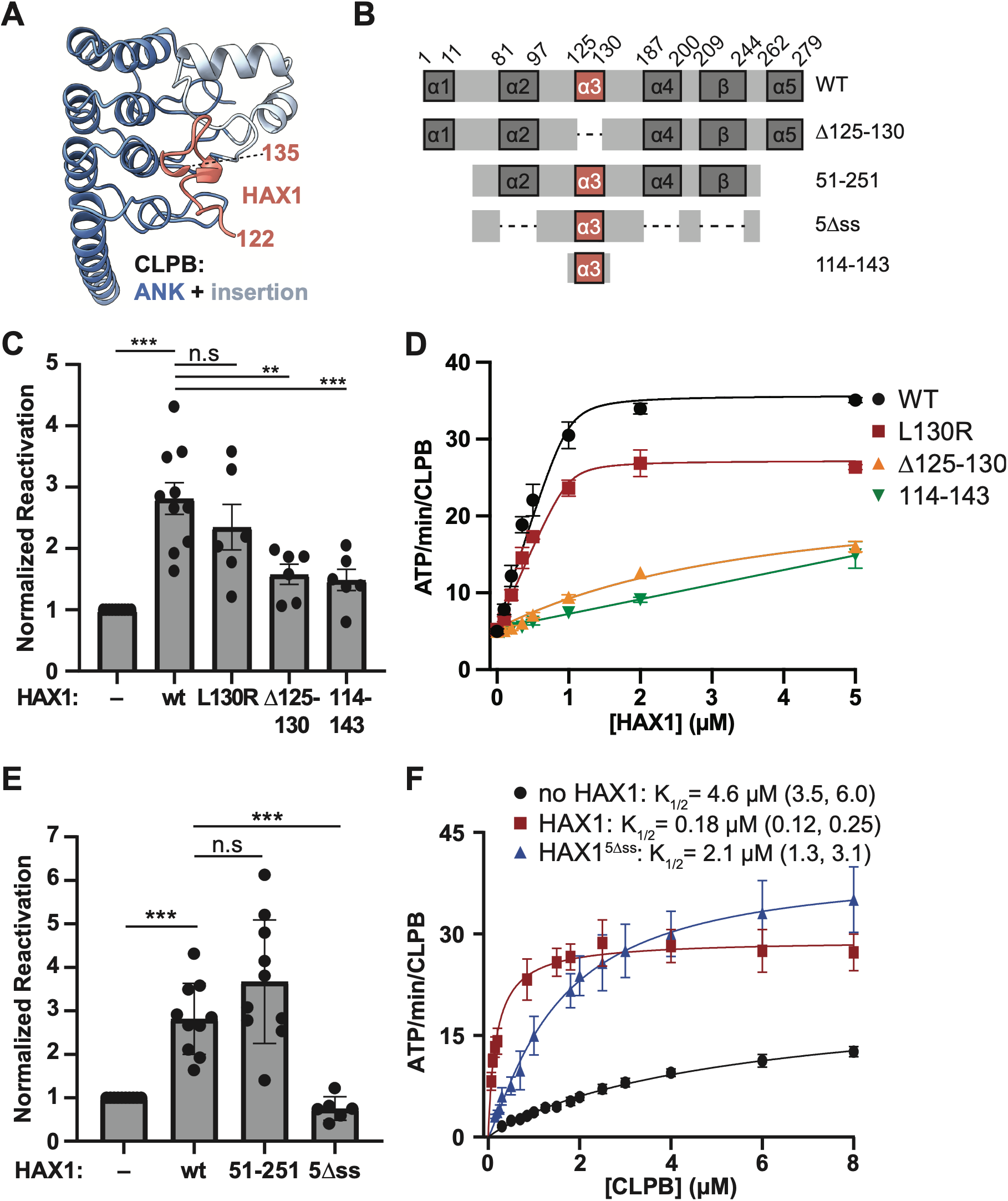
A short helix of HAX1 is necessary for CLPB binding but is insufficient to stimulate CLPB activities. (A) Prediction of the CLPB-HAX1 interaction site, using AlphaFold3. HAX1 is depicted in salmon and the CLPB ankyrin repeat domain is depicted in blue, with the linked subdomain in light blue. (B) Map of HAX1 and truncated variants, with predicted secondary structures indicated. (C) Relative reactivation of aggregated firefly luciferase of 0.25 μM CLPB ± 0.25 μM HAX1 (WT or variants in predicted CLPB-interaction region), n = 10 for wildtype and 6 for variants. (D) ATP hydrolysis rate of 1 μM CLPB in presence of 0-5 μM HAX1 variants, n = 2. p < 0.0001 for each variant differing from wildtype HAX1. (E) Reactivation of firefly luciferase aggregates by 0.25 μM CLPB in the presence of HAX1 variants lacking several elements of predicted secondary structure, n = 10 for wildtype and 6 for variants. (F) Oligomerization of CLPB, monitored using ATP hydrolysis as a proxy. HAX1 variants, when present, are included at a 2:1 molar ratio with CLPB, n = 22 for CLPB alone, n = 10 for CLPB with HAX1, n = 7 for CLPB with HAX1^5Δss^. The p value for K_1/2_ between HAX1 and HAX1^5Δss^ = 0.0033; the p value for k_cat_ = 0.032. For all plots, error = SEM; 95% confidence interval indicated for parameters extracted from fitted equations. Statistical significance denoted by p < 0.05 (*), p < 0.005 (**), p < 0.0005 (***).

We then asked whether the predicted HAX1^125-130^ contact with CLPB was sufficient to stimulate CLPB, using an MBP fusion of a 30-residue peptide from HAX1 centered on this contact (HAX1^114-143^). Although CLPB binding was equivalent to full-length HAX1, as assessed by co-isolation (Fig. S2A), HAX1^114-143^ did not stimulate CLPB disaggregase activity nor specifically stimulate ATPase activity (Fig. 2C,D). Therefore, although the α3 region of HAX1 is sufficient for CLPB binding, it is insufficient for activation. HAX1 contains six regions of predicted secondary structure, including α3 (Fig. 2B). We truncated each of the other five predicted regions to determine how they contribute to stimulating CLPB activity. Individual and paired truncations of HAX1 secondary structures had only small effects on activation of CLPB by HAX1 (Fig S2B-D). A HAX1 variant preserving only the α3 helix, HAX1^5Δss^, still interacted with CLPB and stimulated its ATPase activity, but HAX1^5Δss^ no longer stimulated the disaggregase activity of CLPB and stimulated its K_app_ for oligomerization much less potently (Fig. 2E,F, Fig. S2E). In size-exclusion chromatography, HAX1 and HAX1^51-251^ elute earlier than expected for their monomeric molecular weight, at a volume similar to that expected for a dimer. HAX1^5Δss^ and HAX1^114-143^ eluted as expected of their monomer size (Fig. S2F). The loss in ATPase and disaggregase stimulation by the variants that do not form dimer-like species suggest that the oligomeric form of HAX1 may contribute to its stimulation of CLPB, perhaps by templating oligomerization of CLPB.

### HAX1 selectively activates CLPB isoform 2

Our model of the CLPB-HAX1 contact, supported by our mutagenesis of HAX1 and prior documented interaction of CLPB ANK domain and HAX1 (Fan et al., 2022), indicates that HAX1 binds a groove in the CLPB ANK domain formed by ankyrin motifs 3 and 4 and a structured element linking motifs 2 and 3 (Fig. 2A). The linker between ankyrin motifs 2 and 3 is extended by thirty residues encoded by exon 5 in the longest isoform of CLPB (isoform 1, here referred to as CLPB^L^) (Fig. 3A). Crystal structures of the ANK domains of both isoforms showed that the CLPB^L^ linker element is fold-switched and positions the additional element encoded by exon 5 into the modeled HAX1-binding site (Fig. 3B). Isoform 2 of CLPB appears vastly predominant in most tissues, but CLPB^L^ (distinguished by the addition of exon 5) appears predominant in testis, suggesting it may have a developmentally specialized role (Fig. 3C) (GTEx, 2025). CLPB^L^ has also been reported to interact more weakly than CLPB with HAX1 in cells (Fan et al., 2022). We thus tested the ability of HAX1 to act on CLPB^L^.

**Figure 3.**
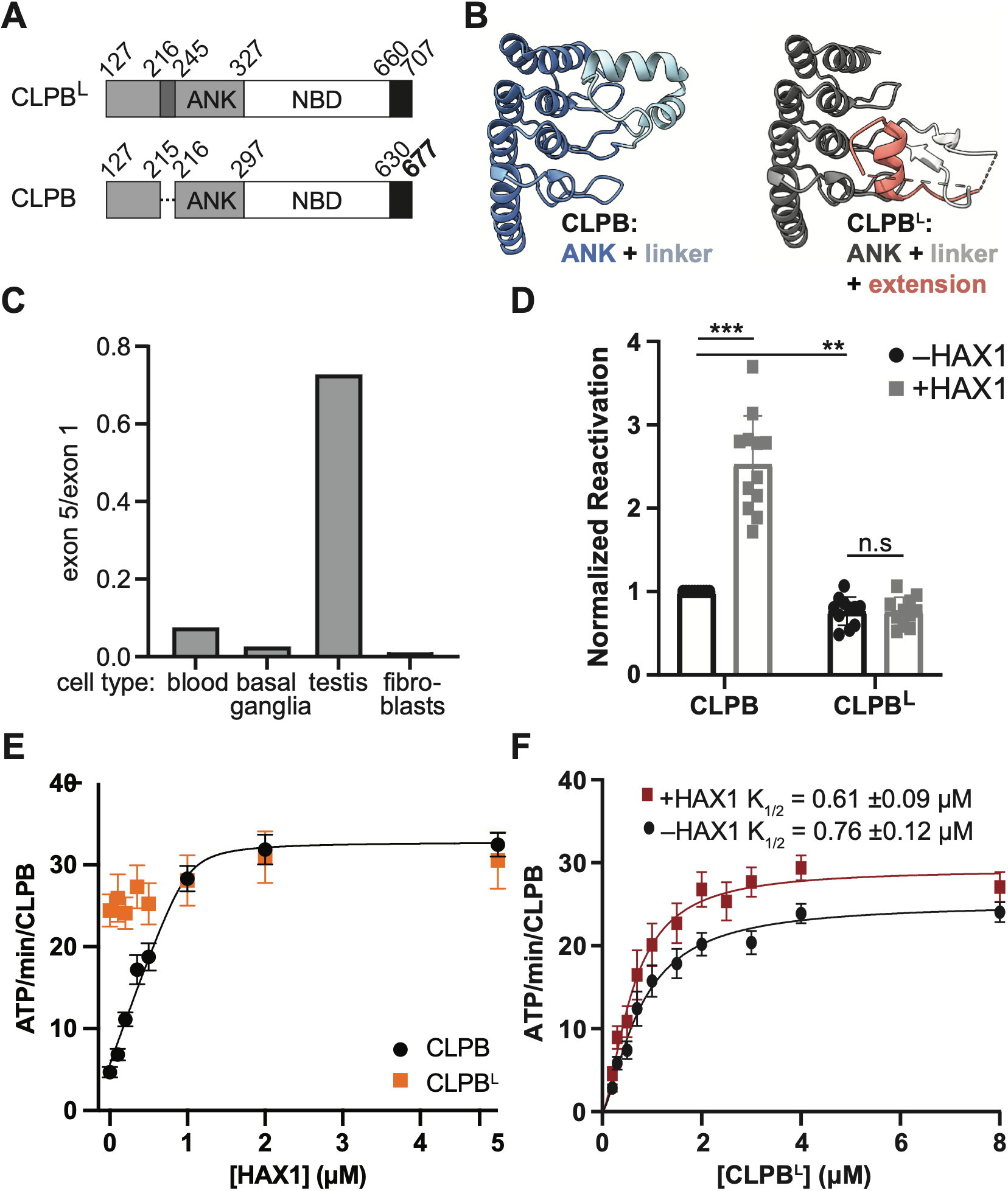
HAX1 specifically stimulates the mature form of CLPB isoform 2. (A) Domain maps of CLPB splice forms (B) Comparison of the ankyrin domain conformation in CLPB (PDB: 8FDS) and CLPB^L^ (PDB: 8DEH). (C) Ratio of expression of exon 1, which is contained in both CLPB and CLPB^L^, and exon 5, which is unique to CLPB^L^, in several representative tissues. Exon expression values were obtained from GTEx Analysis Release V10. (D) Relative reactivation of aggregated firefly luciferase by 0.25 μM CLPB isoforms and cleavage variant ± 0.25 μM HAX1 WT, n = 10. (E) ATP hydrolysis rate of 1 μM CLPB isoforms and cleavage variant in presence of 0-5μM HAX1 WT, n for CLPB = 12, for CLPB^L^ = 6. (F) Oligomerization of CLPB^L^, monitored using ATP hydrolysis rate as a proxy. HAX1, when present, is included at a 2:1 molar ratio with CLPB, n = 18 without HAX1 and n = 14 with HAX1. Data were fit to an allosteric sigmoidal equation. The p value for K_1/2_ = 0.13. For all plots, error = SEM; 95% confidence interval indicated for parameters extracted from fitted equations. Statistical significance denoted by p < 0.05 (*), p < 0.005 (**), p < 0.0005 (***).

HAX1 failed to stimulate the disaggregase activity of CLPB^L^ and only slightly stimulated its ATPase activity (Fig. 3D, E). We observed that CLPB^L^ exhibited an order of magnitude higher-affinity oligomerization than CLPB (K_1/2_ of 0.61 μM for CLPB^L^, compared to 4.6 μM for CLPB) (Fig. 1C, Fig. 3F), but the apparent affinity of oligomerization was not further increased by HAX1. CLPB^L^ also interacted more weakly with HAX1 than CLPB, most dramatically for HAX1^114-143^, the region of HAX1 that our model indicates occupies the same site as the unique element of CLPB^L^ (Fig. 3B, Fig S3E). These data indicate that the altered ANK architecture of CLPB^L^ blocks HAX1 activity but also increases the affinity of CLPB^L^ for oligomerization. These data suggest that the additional element encoded in exon 5 of CLPB^L^ may partially replace HAX1 in function.

Two recessive mutations in CLPB that are linked to MGCA7 lie in ankyrin motif 3, proximal to the site where the exon-5-encoded element and HAX1 dock (T268M and A269T in CLPB^L^, T238M and A239T in CLPB) (Kiykim et al., 2016; Saunders et al., 2015). These mutations partially impair disaggregase activity of CLPB^L^ (Cupo and Shorter, 2020; Lee et al., 2023). We therefore tested how these mutations affected the activity of CLPB, which lacks the proximal exon-5-encoded element of CLPB^L^, and how they affected HAX1 activation of CLPB. Both MGCA7 mutations impaired the disaggregase activity of CLPB alone (monitored at a concentration at which self-assembly is efficient) (Fig. S3B). HAX1 interacted with and stimulated both the ATPase and disaggregase activities of CLPB MGCA7 variants similarly to wildtype CLPB (Fig. S3A,C,E). These data therefore indicate that these MGCA7 mutations within the ANK domain are deleterious independently from the function of either ANK-interacting element, the exon 5 product or HAX1.

### HAX1 enhances client refolding by CLPB

Human CLPB recently was demonstrated to assist in refolding of client proteins (Gupta et al., 2023). Although this activity was directly demonstrated with a soluble unfolded client, it may also contribute to the refolding of clients after disaggregation (thus contributing to measurements of reactivation, as for aggregated luciferase). This activity has been proposed to rely on sequestration of denatured clients in the multicellular-specific enclosure formed by the ANK domains within the CLPB dodecamer (Gupta et al., 2023). We next asked how HAX1 affects CLPB refoldase activity, monitoring the activity of denatured soluble luciferase after dilution into nondenaturing conditions. Without a chaperone, denatured luciferase (100 nM) refolded into its active form only minimally (plateauing at ∼10%, Fig. 4A). CLPB (0.5 μM) increased luciferase refolding, but addition of equimolar HAX1 increased the rate and yield of refolded, active luciferase (Fig. 4A). HAX1 alone did not stimulate refolding (Fig. 4A, E). We additionally observed that CLPB refoldase activity requires ATP hydrolysis; a hydrolysis-blocking mutation in the Walker B motif of CLPB (E425Q) abolished stimulation of refolding (Fig. S4B,C). To stimulate refolding by CLPB, HAX1 required its α3 contact site as well as additional predicted secondary structure; neither HAX1^Δ125-130^, HAX1^114-143^, nor HAX1^5Δss^ stimulated luciferase refolding with CLPB (Fig. 4B,C). This stimulation was specific to HAX1, not a generic feature; MBP did not increase refolding by CLPB and casein slightly inhibited CLPB-stimulated refolding (Fig S4A). Refolding by CLPB^L^ was unaffected by HAX1 and refolding by MGCA7 variants of CLPB was stimulated by HAX1 similiarly to that of wildtype CLPB (Fig. 4D, S4B). Therefore, HAX1 requires the same features to accelerate refolding by CLPB as it does for the combined disaggregation and refolding of an aggregated substrate.

**Figure 4.**
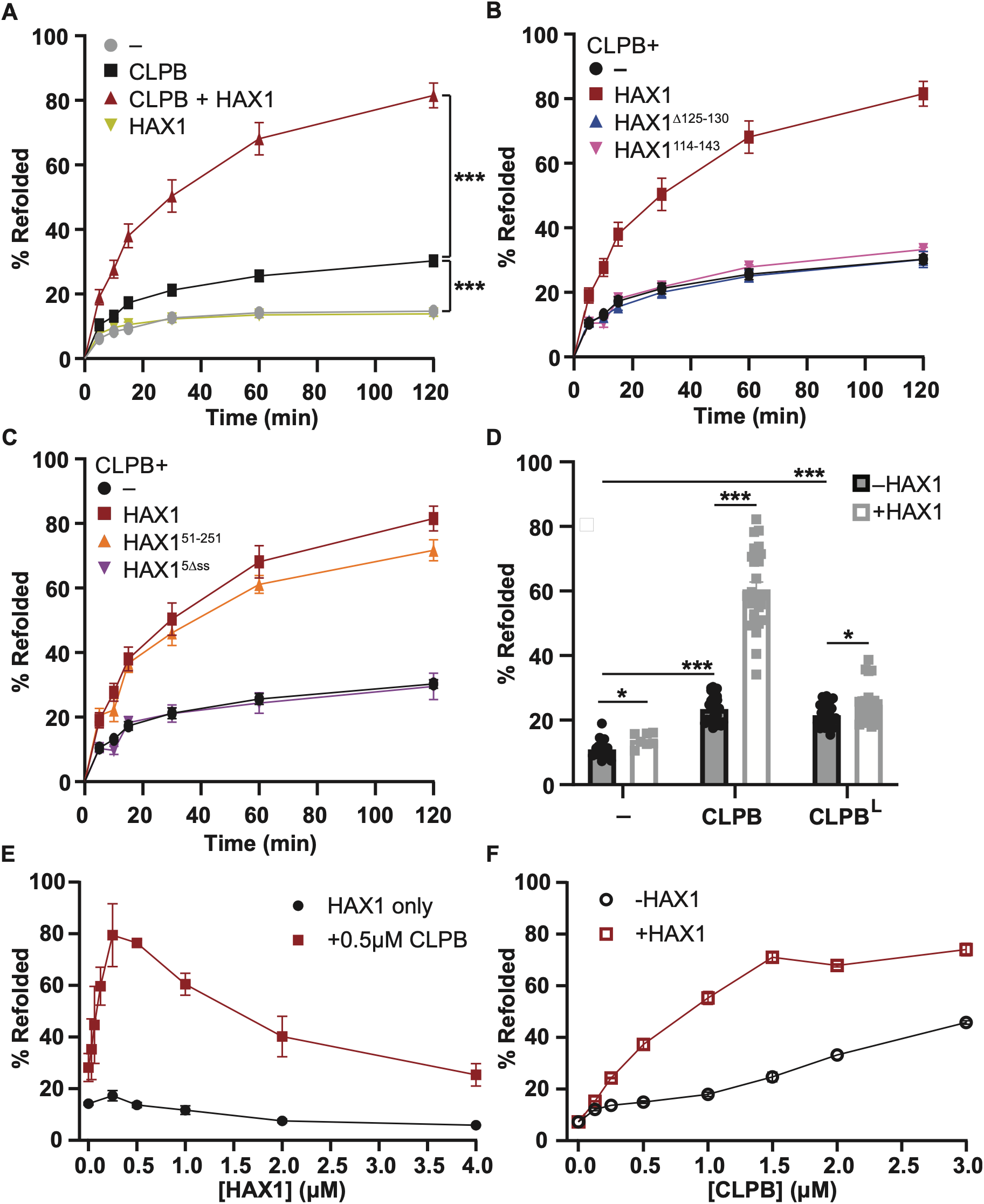
CLPB-HAX1 complexes have enhanced refoldase activity compared to CLPB homo-oligomers. (A) Refolding of soluble denatured firefly luciferase, monitored by luciferase activity over time, with 0.5 μM CLPB ± 0.5 μM HAX1 or HAX1 alone as indicated, n ≥ 4. P-values compare -CLPB vs +CLPB and CLPB vs CLPB+HAX1 refolding after 60 min. (B, C) The effect of 0.5 μM HAX1 variants with mutations in (B) the predicted CLPB interaction site or (C) the predicted secondary structures on 0.5 μM CLPB refoldase activity compared to HAX1 WT, n ≥ 4 (D) Refoldase activity of 0.5 μM CLPB isoforms ± 0.5 μM HAX1 WT, n ≥ 7. (E) The effect of increasing HAX1 concentration (0-4 μM) on the refoldase activity of 0.5 μM CLPB, n ≥ 5. (F) The effect of equimolar HAX1 on increasing CLPB concentration (0.125-3.0 μM) on the fraction of luciferase refolded after 15 minutes, n ≥ 6. Denatured luciferase was diluted to 100 nM in assay buffer. For all plots, error = SEM. Statistical significance denoted by p < 0.05 (*), p < 0.005 (**), p < 0.0005 (***).

We previously observed that CLPB-HAX1 complexes at higher concentrations exhibited reduced disaggregase activity relative to CLPB alone, which we propose could be due to competition between disordered regions of HAX1 and client in trans at another CLPB oligomer. We also infer that HAX1 stimulation of disaggregase is less potent at higher CLPB concentrations because CLPB oligomerization is less dependent on HAX1 at these concentrations. We therefore sought to test how the stimulation of CLPB refoldase by HAX1 was affected by CLPB and HAX1 concentration. To test the stoichiometry at which HAX1 and CLPB operate as a refoldase, we titrated HAX1 against 0.5 μM CLPB and monitored activation of denatured luciferase. We observed that CLPB refoldase activity was maximal at approximately equimolar HAX1, similarly to its disaggregase activity. Superstoichiometric HAX1 inhibited refolding of luciferase by CLPB-HAX1, consistent with its possible competition for engagement by CLPB (Fig. 4E, S4D). We also compared the refoldase activity of 1:1 CLPB-HAX1 or CLPB alone at higher concentrations (Fig. 4F). We observed that CLPB-HAX1 sustained enhanced refoldase activity relative to CLPB alone at all concentrations monitored (up to 3 μM). CLPB-HAX1 exhibited a dose-dependent increase in refolding up to 1.5 μM. Above this concentration, CLPB oligomers (hexameric or dodecameric) are in excess of the denatured substrate (100 nM), suggesting that CLPB-HAX1 activity does not increase further because all substrates are engaged. Because of limitations of denatured luciferase solubility and denaturant concentration in the assay, we were not able to assess refolding of luciferase at higher concentrations, and so could not fully separate the ability of HAX1 to stimulate CLPB refolding from its stimulation of oligomerization. However, these data indicate that CLPB-HAX1 complexes are potent in their ability to refold a denatured substrate.

### HAX1 and CLPB form a stoichiometric oligomer containing sub-dodecameric CLPB

We infer from the peak in disaggregase (Fig. 1D) and refoldase (Fig. 4E) activity at or close to 1:1 HAX1:CLPB that they likely operate most efficiently as a 1:1 complex, but we sought a physical measurement of this stoichiometry as well as information about the oligomeric state of this complex. In addition to a hexamer, CLPB can form a dodecamer from two hexamers mediated by cross-hexamer ANK domain contacts. Because of the lower resolution of cryoEM maps of the ANK-ANK interface, the precise contacts mediating this interface were not visible (Cupo et al., 2022; Gupta et al., 2023; Wu et al., 2023), but because HAX1 binds in this region, it could either stabilize or inhibit dodecamer formation.

To monitor the oligomeric state of HAX1 and CLPB, we used size-exclusion chromatography. To stabilize CLPB oligomers for these observations, we used CLPB^E425Q^, which harbors a mutation in the Walker B domain that allows binding but not hydrolysis of ATP. This variant of CLPB migrated primarily as an apparent dodecamer, consistent with previous observations of isoform 1 of CLPB in nucleotide-bound, non-hydrolyzing conditions (Gupta et al., 2023). To observe CLPB-HAX1 complexes, we mixed CLPB^E425Q^ with excess HAX1^51-251^. This HAX1 variant retained the ability to activate CLPB (Fig. 2E, Fig. S2E) but had a more defined elution profile than full-length HAX1 (Fig. S2F), simplifying interpretation of its effect on CLPB oligomerization. Addition of HAX1^51-251^ shifted the CLPB peak to form a left shoulder and depleted the HAX1-only peak, indicating formation of a slightly larger CLPB-HAX1 complex (Fig. 5A). Quantitation of relative amounts of CLPB and HAX1 across the peak a nearly stoichiometric ratio (0.7-1) of CLPB/HAX1 molecules (Fig. 5E). Because HAX1^51-251^ and CLPB are similar in size (67 and 63 kDa, respectively), this minor shift we observe is not consistent with binding of 8-12 molecules of HAX1 to a CLPB dodecamer but instead indicates that HAX1 binding induces formation of a lower-order oligomer of CLPB (Fig. 5F). HAX1^Δ125-130^, which lacks the primary CLPB-interacting site, co-migrates less efficiently and does not induce a similar shift in CLPB migration (Fig. S5).

**Figure 5.**
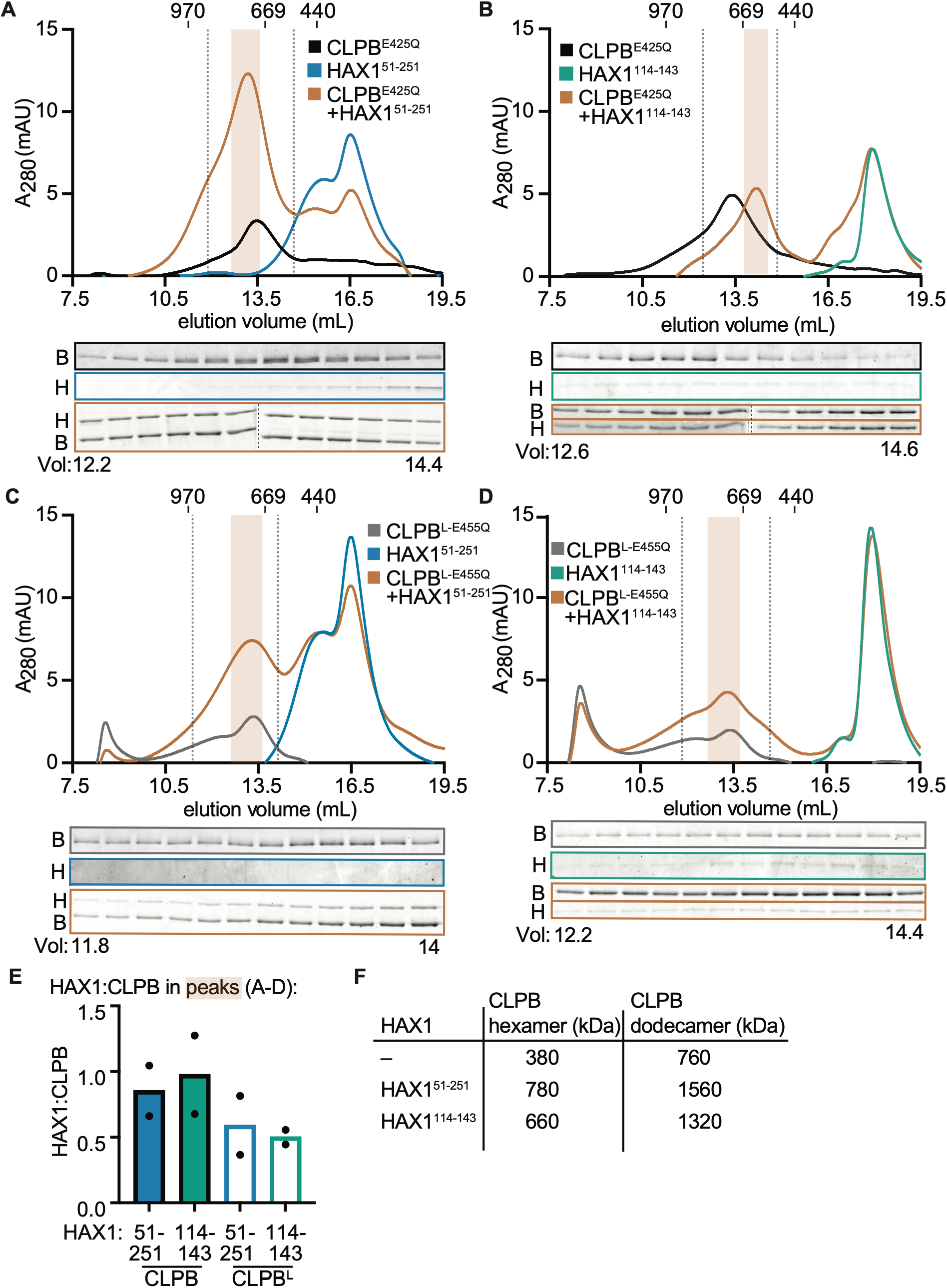
HAX1 induces formation of a smaller oligomer of CLPB. The size of CLPB, HAX1, and CLPB-HAX1 complexes were assessed using size exclusion-chromatography. Samples were prepared with an ATP-hydrolysis-blocked Walker B variant of CLPB (E425Q for isoform 2 or E455Q for isoform 1) (32 μM) and/or HAX1 (37.5 μM, in slight excess) in the presence of 2 mM ATP and separated on a Superose 6 Increase 10/300 column, equilibrated in the same buffer with 2 mM ATP. Elution fractions highlighted between vertical dashed lines were analyzed by SDS-PAGE and Sypro Red or Orange staining. At the top of each chromatogram, the molecular weight (in kDa) of three protein standards is noted at their elution volumes. (A) CLPB^E425Q^ and HAX1^51-251^; (B) CLPB^E425Q^ and HAX1^114-143^. (C) CLPB^L-E455Q^ and HAX1^51-251^; (D) CLPB^L-E455Q^ and HAX1^114-143^. (E) The ratios of HAX1:CLPB molecules were by quantifying the fluorescence intensity of protein bands in SDS-PAGE gels from size exclusion eluate fractions in comparison with concentration standards of these proteins. The average HAX1:CLPB across the peak of interest highlighted in orange on each chromatogram is plotted for each CLPB-HAX1 combination. (F) Molecular weights of possible CLPB homo-oligomer and CLPB-HAX1 stoichiometric oligomer complex formations. For all chromatograms n = 2.

We tested whether HAX1-induced remodeling of CLPB dodecamers is recapitulated by HAX1^114-143^, which retained the ability to bind CLPB while failing to stimulate its activity (Fig. 2C,D). Addition of HAX1^114-143^ shifted the elution profile of CLPB^E425Q^ towards a larger elution volume, revealing the formation of an unambiguously smaller CLPB-HAX1 complex that also contains approximately stoichiometric CLPB/HAX1 (Fig. 5B,E,F). Because HAX1^114-143^ fails to activate CLPB (Fig. 2C,D), this shift demonstrates that induction of a smaller oligomer of CLPB is not sufficient for activation.

We also asked how CLPB-HAX1 oligomer formation may differ with CLPB^L^, which is not stimulated by HAX1 (Fig. 3). The analogous Walker B variant of CLPB^L^, CLPB^L-E455Q^, migrated similarly to CLPB, eluting at a volume consistent with a dodecamer (Fig. 5C,D). Addition of HAX1^51-251^ caused a smaller shift of the CLPB^L^ peak and less depletion of the HAX1-alone peak than we observed for CLPB (Fig.5C). Some HAX1 did co-migrate with CLPB, but at a lower ratio of 0.4-0.8 HAX1:CLPB (Fig. 5C,E). Similarly, addition of HAX1^114-143^ did not significantly shift the elution of CLPB^L^, and the CLPB^L^ peak contained (Fig. 5D). The peak eluting from this mixture contained a ratio of about 0.4-0.6 HAX1:CLPB (Fig. 5E). These results support the conclusion that HAX1 binding and complex formation is less efficient with CLPB^L^ compared to CLPB and additionally indicate that HAX1 does not induce a dramatic change in the oligomer that CLPB^L^ forms, as it does for CLPB.

## DISCUSSION

Previous genetic, proteomic, and phenotypic links between CLPB and HAX1 (Baker et al., 2024; Chen et al., 2019; DepMap and Broad, 2025; Fan et al., 2022; Wakula et al., 2020; Wu et al., 2023), as well as the closely overlapping syndromes that mutations in these two genes cause (Klein et al., 2007; Saunders et al., 2015; Warren et al., 2022; Wortmann et al., 2015), indicated that their functions were closely linked. The biochemical investigation we present here establishes a mechanism for this link: HAX1 is an activating cofactor of CLPB. HAX1 strengthens the oligomerization of CLPB into an active complex by more than an order of magnitude, thus promoting its activity as a disaggregase and as a refoldase. To do this, a short helical region in HAX1 (α3) is required to form a contact with the ANK domain of CLPB, but this contact is not sufficient to promote CLPB activity. Additional predicted secondary structural elements in HAX1 are required for this binding to activate CLPB, either through interaction between HAX1 molecules within a CLPB oligomer, through an additional weak contact with CLPB, or by an allosteric effect.

Our data indicate that HAX1 activation is most potent when it is near-stoichiometric with CLPB. We additionally observe that HAX1 binding may break the CLPB dodecamer into a smaller hetero-oligomer whose apparent size and stoichiometry is most consistent with a hexamer of CLPB, elaborated with up to six HAX1 molecules. Based on the position of HAX1 contact on the ANK domain of CLPB, HAX1 insertion at this site may either directly block ANK-ANK contacts that form the dodecamer or may alter the conformation or positioning of the ANK domain to disfavor dodecamer formation. We observe CLPB-HAX1 complexes to be active both in reactivation of an aggregated client and in refolding of a denatured client, suggesting that the enclosed ANK domain cage of the dodecamer may not be required for client refolding by CLPB-HAX1. Alternatively, the apparent hexamer of CLPB elaborated with HAX1 we observe with ATP-locked CLPB may be only one of several states that an active complex cycles through during substrate renaturation.

The activation of CLPB by HAX1 has conceptual similarity to the activation of unicellular ClpB homologs by Hsp70. Of the two AAA+ domains in unicellular ClpB homologs, the C-terminal AAA+ domain (AAA2) provides the primary force for protein unfolding (Deville et al., 2019); this is the domain conserved in metazoan ClpB homologs. The activity of unicellular ClpB homologs is constitutively repressed by interactions between a regulatory coiled-coil middle (M) domain and the N-terminal AAA+ (AAA1) domain. This inhibition is relieved by contact with Hsp70 engaged with aggregated proteins, activating ClpB for protein disaggregation (Deville et al., 2019; Lee et al., 2013; Oguchi et al., 2012; Rosenzweig et al., 2013; Seyffer et al., 2012). In metazoan ClpB homologs, the regulatory module thus formed by the AAA1 and M domains of unicellular ClpB has been replaced by an ankyrin-repeat domain, which we here show makes activating contact with the intrinsically disordered protein HAX1. By potentiating CLPB activity, HAX1 could control CLPB as Hsp70 does unicellular ClpB, but through a distinct mechanism. We find that several conserved elements of HAX1 are dispensable for activation of CLPB in vitro; these elements might direct high-priority clients to CLPB or coordinate its activity with other proteostasis machinery in the IMS.

## MATERIALS AND METHODS

### Plasmid construction

CLPB constructs (CLPB, CLPB^L^, and Walker B variants) were cloned with an N-terminal H_10_-SUMO tag into pET28b vector with kanamycin resistance. AgeI and XhoI sites were used to insert CLPB^L^ or CLPB sequence. Walker B variants were produced using the wildtype template and mutagenic primers. All H_6_-tev-MBP tagged HAX1 constructs were cloned into pET23a. HAX1 sequence was inserted using BamHI and XhoI restriction enzyme sites. HAX1 variants were produced using wildtype template and mutagenic primers. Untagged HAX1 and H_6_-tev-FFL were cloned into pET28b.

### CLPB expression and purification

CLPB constructs were transformed into BL21 Rosetta cells and grown on LB-Agar plates containing 30 μg/ml kanamycin and 20 μg/ml chloramphenicol. Starter cultures were inoculated overnight at 37°C and diluted into 1L cultures at 1:100 dilution. Cultures were induced with 0.5mM IPTG at 16°C for 15-18 hours after attaining an optical density of 0.5-0.7. Cell pellets were then harvested at 5000 x *g*, washed and stored in -80°C until ready to use.

CLPB pellets were resuspended in lysis buffer (500 mM NaCl, 10% glycerol, 20 mM imidazole, 50 mM Tris pH 7.8, 1 mM DTT, 0.5 mM PMSF) and passed three times through an LM20 microfluidizer (Microfluidics) at 18,000 psi. The lysate was then clarified at 13,000 x *g* for 20 min. Then the supernatant was poured into a pre-equilibrated column containing Ni-NTA resin and allowed to bind. The CLPB-bound resin was then washed extensively with lysis buffer, followed by an intermediate wash step with lysis buffer containing 50 mM imidazole, and eluted in buffer containing 250 mM imidazole. For CLPB (CLPB isoform 2), protein elutions were pooled, treated with Ulp1 to remove the SUMO-tag and dialyzed in 10kDa cutoff bag against storage buffer SA (500 mM NaCl, 10% glycerol, 25 mM Tris pH 8.5, 1 mM DTT). CLPB^L^ (CLPB isoform 1) variants were dialysed against storage buffer SB (500 mM NaCl, 10% glycerol, 25 mM Tris pH 7.8, 1 mM DTT). Thereafter, proteins were concentrated in 30 kDa cutoff concentrators (Sartorius). Concentrated proteins were injected into a Superose 6 Increase 10/300 GL column equilibrated in their respective storage buffers. Fractions were pooled, supplemented with glycerol to 20%, concentrated, flash frozen, and stored at -80°C.

For Walker-B variant CLPB used in analytical SEC experiments, during 50 mM imidazole washes, 5 mM ATP and 5 mM MgCl_2_ was added to release contaminating bacterial chaperone proteins.

### HAX1 expression and purification

H_6_-tev-MBP-HAX1 was expressed in BL21 Rosetta cells in the presence of kanamycin (30 μg/ml) and chloramphenicol (20 μg/ml), induced at 0.5-0.7 OD with 0.5 mM IPTG for 15-18 hours at 16°C. Pellets were harvested, washed in PBS, re-pelleted and stored in either -20 or -80°C. H_6_-tev-MBP-HAX1 pellets were lysed in buffer A (250 mM NaCl, 5% glycerol, 20 mM imidazole, 25 mM Tris pH 7.8, 1 mM DTT, 0.5 mM PMSF) at 18,000 psi through a microfluidizer. The clarified lysate was allowed to bind to pre-equilibrated Ni-NTA resin by gravity flow. The resin was then washed with 10x CV buffer A. Proteins were eluted in buffer C (250 mM NaCl, 5% glycerol, 250 mM imidazole, 25 mM Tris pH 7.8, 1 mM DTT). Pure fractions were pooled and dialysed in 10 kDa cutoff bag in storage buffer SD (250 mM NaCl, 5% glycerol, 25 mM Tris pH 7.8, 1 mM DTT). After dialysis, proteins were concentrated in 30 kDa cutoff concentrators. Concentrated proteins were injected into a Superose 6 increase 10/300 GL column equilibrated in storage buffer SD. The pure fractions were pooled, supplemented with a final 15% glycerol content, concentrated, flash frozen in liquid N_2_ and stored at -80°C. Expression and purification protocol was followed for the MBP tag control.

To isolate untagged HAX1 from inclusion bodies, BL21 Rosetta cells containing the construct were grown to OD 0.5-0.7 in LB (Lennox) and induced with 0.5 mM IPTG at 37°C for 3-4 hr. Cells were harvested and pellets stored in -20°C. Pellets were resuspended in lysis buffer (500 mM NaCl, 10% glycerol, 50 mM Tris pH 7.8, 2 mM DTT), microfluidized at 18,000 psi and centrifuged at 5000 x *g*. The supernatant was discarded and the pellet was resuspended in IB wash buffer (1.0M urea, 500 mM NaCl, 10% glycerol, 50 mM Tris pH 7.8, 2 mM DTT). The washed inclusion body was either stored at -20°C or resuspended in solubilization buffer (4.0 M urea, 500 mM NaCl, 10% glycerol, 50 mM Tris pH 7.8, 2 mM DTT). The inclusion body was then homogenized by sonication and then clarified at 13,000 x *g*. The supernatant was then dialyzed in storage buffer SB. The dialysate was then centrifuged to remove insoluble precipitates. The supernatant was then concentrated with 10 kDa cutoff centrifugal devices. The concentrated sample was then clarified by centrifugation, flash frozen and stored at - 80°C.

### Firefly luciferase expression and purification

H_6_-tev-firefly luciferase was expressed in Rosetta cells induced at with 0.5 mM IPTG for 15-18 hours at 16°C. Pellets were harvested, washed in PBS, re-pelleted and stored in - 80°C. FFL pellets were lysed in buffer A (see HAX1 expression for details) at 18,000 psi by microfluidizer. The clarified lysate was allowed to bind to pre-equilibrated Ni-NTA resin by gravity flow. The FFL-bound resin was then washed 5x CV buffer A, 5x CV buffer B (250 mM NaCl, 5% glycerol, 50mM imidazole, 25 mM Tris pH 7.8, 1 mM DTT, 0.5 mM PMSF). FFL proteins were eluted in buffer C. Pure fractions were pooled and dialysed in 10 kDa cutoff bag in storage buffer SD. After dialysis, proteins were concentrated in 30 kDa cutoff cassettes. Concentrated proteins were injected into Superdex 200 increase 10/300 GL column equilibrated in storage buffer SD. The pure fractions were pooled, supplemented with 15% glycerol final, concentrated, flash frozen and stored at -80°C.

### ATPase assay

Proteins were prepared in assay buffer (150 mM KCl, 50 mM HEPES pH 7.2, 8 mM MgCl_2_). CLPB proteins (4 μM) were mixed in a 1:1 volume ratio with increasing concentration of either HAX1 variants or control proteins (fitc-casein, MBP tag, luciferase precipitates). The buffer only, CLPB only, CLPB+protein or protein only mix (10 μl) were pipetted into designated wells of transparent 384-well microplates. The microplate was pre-incubated at 37°C. NADH-coupled ATPase assay components (10 mM NADH, 15 mM phosphoenolpyruvate, 30 U/mL pyruvate kinase, 40 U/mL lactate dehydrogenase and 2 mM ATP) was also incubated at 37°C for 3 min. To begin the reaction, 10 μl of the NADH-enzyme-ATP mix was added to the wells containing CLPB, HAX1, protein controls or buffer. The NADH absorbance at 340 nm was monitored at time intervals on SpectraMax M5^e^ plate reader. The rate of NADH absorbance loss was extracted from SoftMax Pro 7.0.3 software and converted into ATP hydrolyzed per minute per CLPB molecule after the blank rates were subtracted. Luciferase precipitates were generated by heating luciferase (20 μM) at 42°C for 15 min.

For CLPB-concentration-dependent ATPase activity, various concentrations of CLPB were prepared and mixed with either buffer or a 2-fold molar excess concentration of HAX1 compared to the CLPB concentration. Similarly, the CLPB-buffer or CLPB-HAX1 samples were pre-incubated at 37°C in the plate reader, the reactions initiated upon addition of NLPP mastermix and A340 monitored at appropriate time intervals to measure the ATPase rate before the NADH is fully depleted.

To measure CLPB ATPase rate under disaggregase conditions, CLPB (1 μM) was mixed with increasing HAX1 concentration in a 1:1 volume ratio. The final HAX1 concentrations are shown in the plotted data. The NLPPA mix was supplemented with 100 nM FFLagg. The samples and initiation mix were incubated at 30°C for 3 min separately and then mixed to begin measuring ATPase rate in the presence of FFL_agg_.

To measure CLPB ATPase rate under refoldase conditions, CLPB (2 μM) was mixed with an equal volume of HAX1 stocks of varied concentrations. The NLPPA mix and the CLPB, HAX1 samples were incubated at 25°C. Immediately before reaction initiation, 10 μM of denFFL was added to the NLPPA mix to a final concentration of 200 nM. The reactions were incubated at 25°C and NADH absorbance at 340nm measured at regular intervals for 2 hr or when signal plateaued.

### Disaggregase assay

This assay was based on a previously published method (Gupta et al., 2023). To prepare firefly luciferase aggregates (FFL_agg_), 20 μM FFL was denatured in 4 M urea dissolved in assay buffer AB (150 mM KCl, 50 mM HEPES pH 7.2, 8 mM MgCl_2_) for 30 min at 28°C. The denatured FFL was then diluted 100-fold in assay buffer and incubated for another 5 min at 28°C. FFL_agg_ (100 nM) was then supplemented with 8mM ATP and ATP regenerating system (ARS: 7mM creatine phosphate, 14U/ml creatine kinase, Sigma 3755). CLPB (1 μM) was then mixed with equal volume of various concentrations of HAX1 variants or protein controls. The FFL_agg_ was then added to the CLPB containing samples at a 1:1 volume ratio to initiate FFL reactivation. The samples were incubated at 30°C for either 90 min or an aliquot was removed to measure reactivation at a specific time point. To measure the luminescence recovered, each sample was transferred to a white opaque 96-well microplate and mixed with 2x substrate mix (200 mM Tris pH 7.8, 10 mM MgCl_2_, 0.3 mM coenzyme A, 0.4 mM ATP, 0.2 mg/mL D-luciferin, 16 mM DTT) at 25°C. A sample of 50 nM untreated firefly luciferase supplemented with ATP and ARS system served as 100% luciferase activity.

### Refolding assay

Denatured FFL (denFFL) was prepared as previously described (Gupta et al., 2023). Briefly, 10 μM FFL was incubated for 1hr in 5 M guanidium chloride dissolved in buffer AB at 25°C. To initiate refolding at 25°C, the denFFL is diluted 100-fold (100 nM final) into buffer containing 4 mM ATP, ARS (2.5 mM creatine phosphate, 5 U/ml creatine kinase) and either CLPB or HAX1 or both. Final protein concentrations are indicated in the figure. Aliquot samples were removed at regular intervals and mixed with equal volume 2x substrate mix to measure chemiluminescence increase over time.

### Analytical size-exclusion chromatography

To assemble CLPB-HAX1 complexes, ∼32 μM CLPB E425Q, ∼37.5 μM MBP-HAX1 and 2 mM ATP were mixed in assay buffer (150 mM KCl, 50 mM HEPES pH 7.2, 1 mM DTT, 8 mM MgCl_2_). Samples were centrifuged at 21,000 x g for 10 min at 4°C to remove potential aggregates, then incubated at 25°C for 10 min to allow for complex assembly. 25 μL samples were injected onto a Superose 6 Increase 10/300 GL column with a Hamilton syringe. Elutions were collected in 200 μL fractions and analyzed by SDS-PAGE, stained with SYPRO Red or Orange. Gels were imaged on a Cytiva Typhoon RGB. Protein band fluorescence intensity was quantified in ImageQuant using rolling-ball background subtraction. Intensity was converted to molecular quantities by comparison with a standard concentration series of each protein stained with the equivalent SYPRO dye.

### Coprecipitation of CLPB and HAX1

His-tag-based pulldowns were performed with Ni^2+^-charged magbeads (Genscript). 10 μg H_6_-tev-MBP-HAX1 variants were mixed with 10 μg CLPB variants and left to sit at 25°C for 10 min. For every 10 μg H_6_-tagged protein, 2 μl of settled beads were used. The bead slurry was equilibrated in bind-block buffer (150 mM NaCl, 50 mM Tris pH 7.8, 6 mM MgCl_2_, 1 mM DTT, 5% milk). The beads for each sample were then resuspended in 500 μl bind-block buffer and added to the designated CLPB-HAX1 mixture bringing CLPB or HAX1 to 0.3 μM. The bead-protein mixture was incubated with head-over-head mixing at 4°C for 1-2 hr. The beads were washed thrice with 500 μl buffer WB (150 mM NaCl, 50 mM Tris pH 7.8, 6 mM MgCl_2_, 1 mM DTT, 20 mM imidazole) and then eluted in 40 μl buffer EB (150 mM NaCl, 50 mM Tris pH 7.8, 6 mM MgCl_2_, 1 mM DTT, 500 mM imidazole). The elutions were supplemented with 10 μl 5X Laemmli buffer, boiled 95°C for 5 min and then analyzed via SDS-PAGE and Coomassie staining.

### Statistical analysis

Single-value comparisons were made using Student’s t-test and curve fits were compared by extra-sum-of-squares F test using GraphPad Prism.

## ACKNOWLEDGMENTS

We thank Jacquelyn LaVallee, Matt Copeland, and Elle Yung for contributions at early stages of this project and members of the Kardon lab for valuable discussions. This work was supported by National Institutes of Health grant R01GM151332 (J.R.K). M.A.V.F. was supported by The Jane Coffin Childs Memorial Fund for Medical Research. J.G.H. was supported by National Institutes of Health grant T32GM135126 and M.K. was supported by National Institutes of Health grant T32GM139798. The content is solely the responsibility of the authors and does not necessarily represent the official views of the National Institutes of Health.

## AUTHOR CONTRIBUTIONS

**Monifa A V Fahie:** Conceptualization, Methodology, Investigation, Supervision, Writing - Original Draft, Writing - Review & Editing. **Julia G Hoffman**: Methodology, Investigation, Writing - Original Draft, Writing - Review & Editing **Mimi S Kay:** Methodology, Investigation, Writing - Review & Editing **Julia R Kardon:** Conceptualization, Methodology, Supervision, Writing - Original Draft, Writing - Review & Editing, Funding acquisition.

---

**Figure S1 related to Figure 1.**
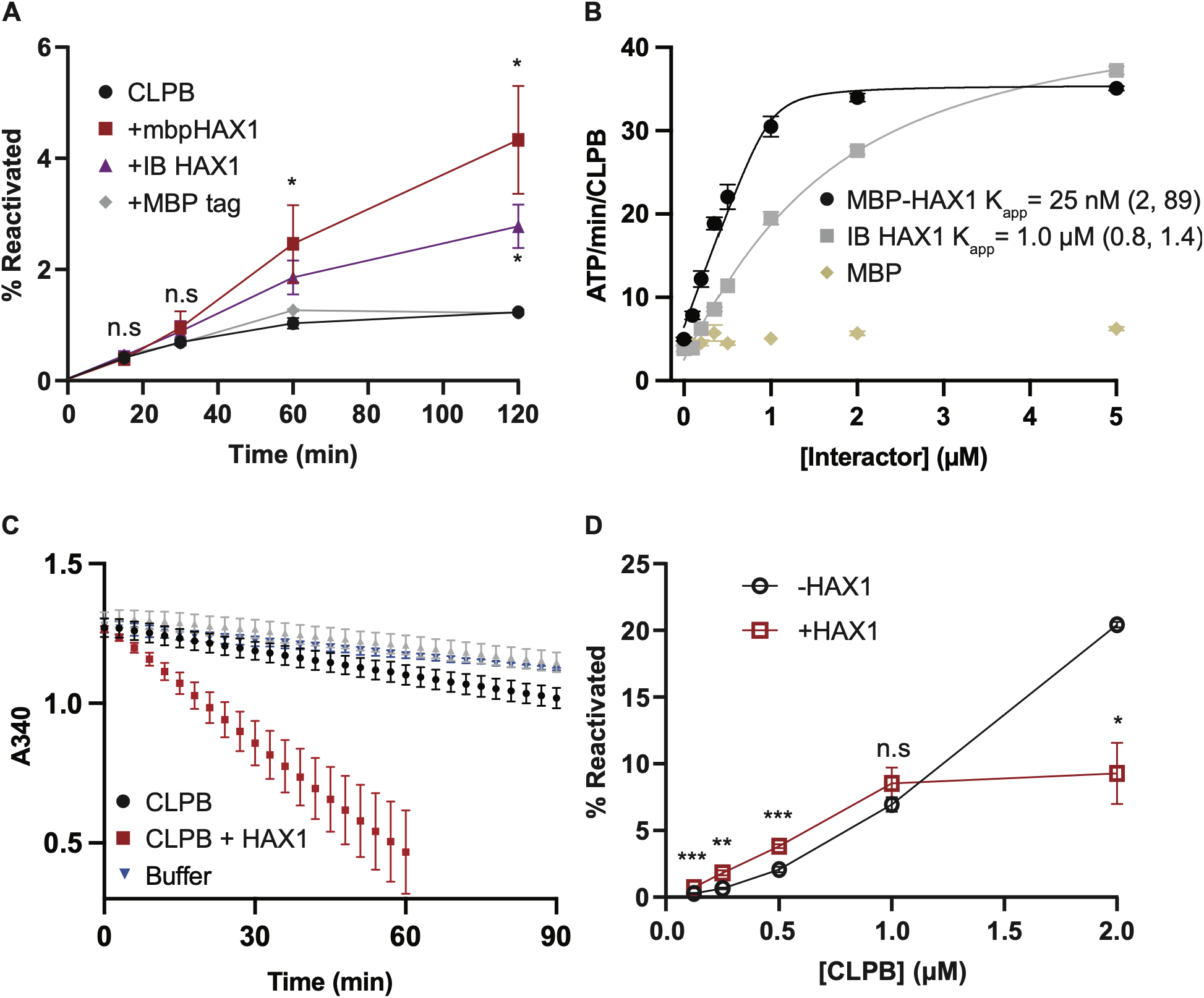
(A) Disaggregase activity of 0.25 μM CLPB ± 0.25 μM MBP-HAX1, untagged HAX1 isolated from inclusion bodies and renatured, and MBP, n = 2. (B) ATPase rate of 1 μM CLPB in the presence of 0-5 μM MBP-HAX1, untagged HAX1, or MBP, n = 2. (C) ATP hydrolysis rate of 0.25 μM CLPB ± 0.25 μM MBP-HAX1 in the presence of 50 nM aggregated firefly luciferase. Assay becomes nonlinear below ∼A_340_ = 0.5 due to depletion of its components. n = 2. (D) Disaggregase activity of 0.125-2.0 μM CLPB in the presence of 0 or equimolar MBP-HAX1, n = 5. For all plots, error = SEM. Statistical significance denoted by p < 0.05 (*), p < 0.005 (**), p < 0.0005 (***).

**Figure S2, related to Figure 2.**
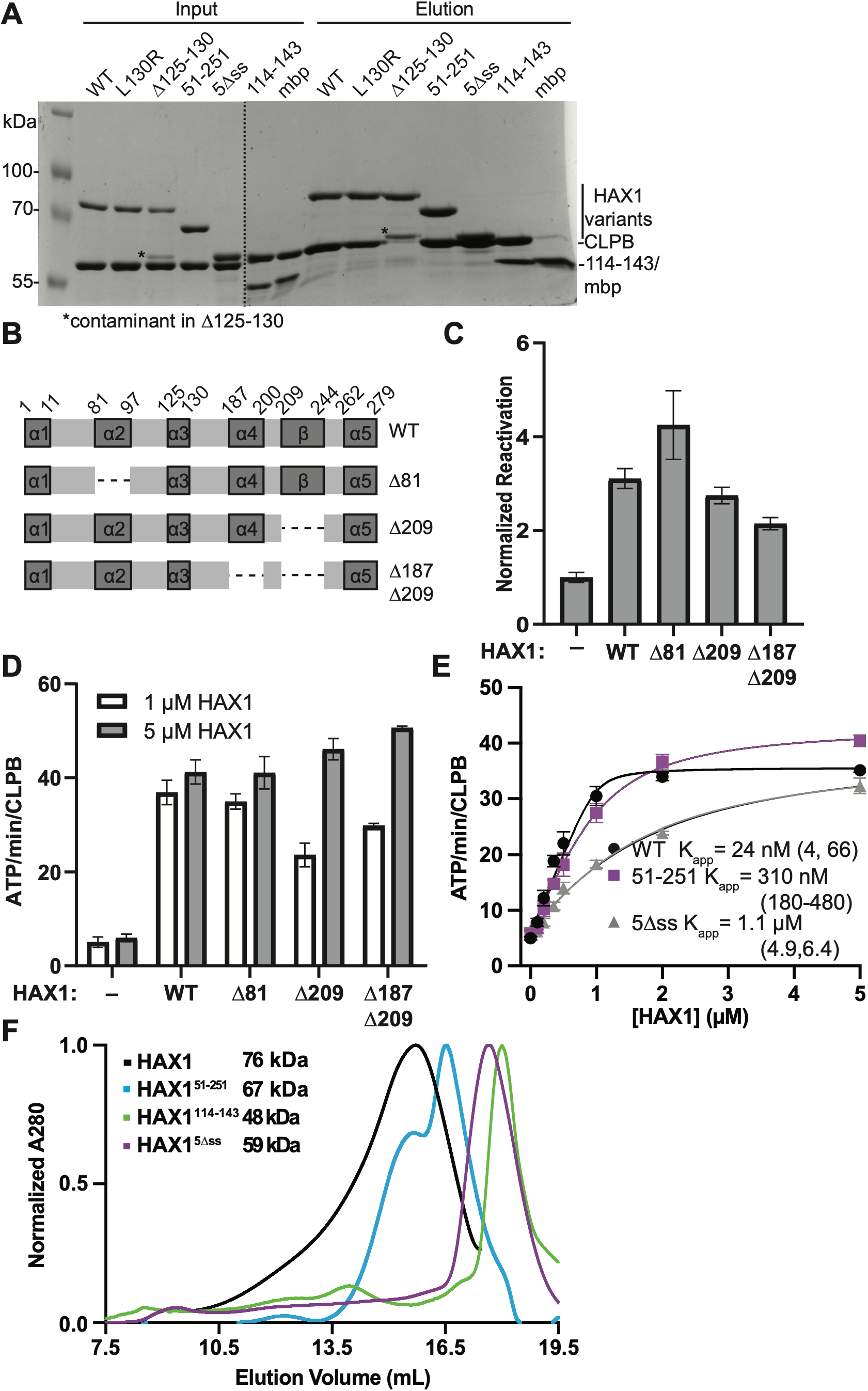
(A) Co-isolation of CLPB with His_6_-tagged HAX1, n = 6. (B) Diagram of HAX1 predicted secondary structure and truncations. (C) Relative reactivation of aggregated firefly luciferase of 0.25 μM CLPB ± 0.25 μM HAX1 variants as indicated, n = 8 for wt and Δ209, n = 6 for Δ81, and n = 4 for Δ187Δ209. (D) k_cat_ for ATP of CLPB with HAX1 variants as indicated, n = 2. (E) ATP hydrolysis activity of 1 μM CLPB in presence of 0-5 μM HAX1 variants, fit to a quadratic velocity equation, n = 2. (F) HAX1 variants were analyzed using a Superose 6 increase 10/300 column, equilibrated in buffer SD (250 mM NaCl, 5% glycerol, 25 mM Tris pH 7.8, 1 mM DTT). For all plots, error = SEM; fitted parameters represented with 95% confidence interval. Statistical significance denoted by p < 0.05 (*), p < 0.005 (**), p < 0.0005 (***).

**Figure S3, related to Figure 3.**
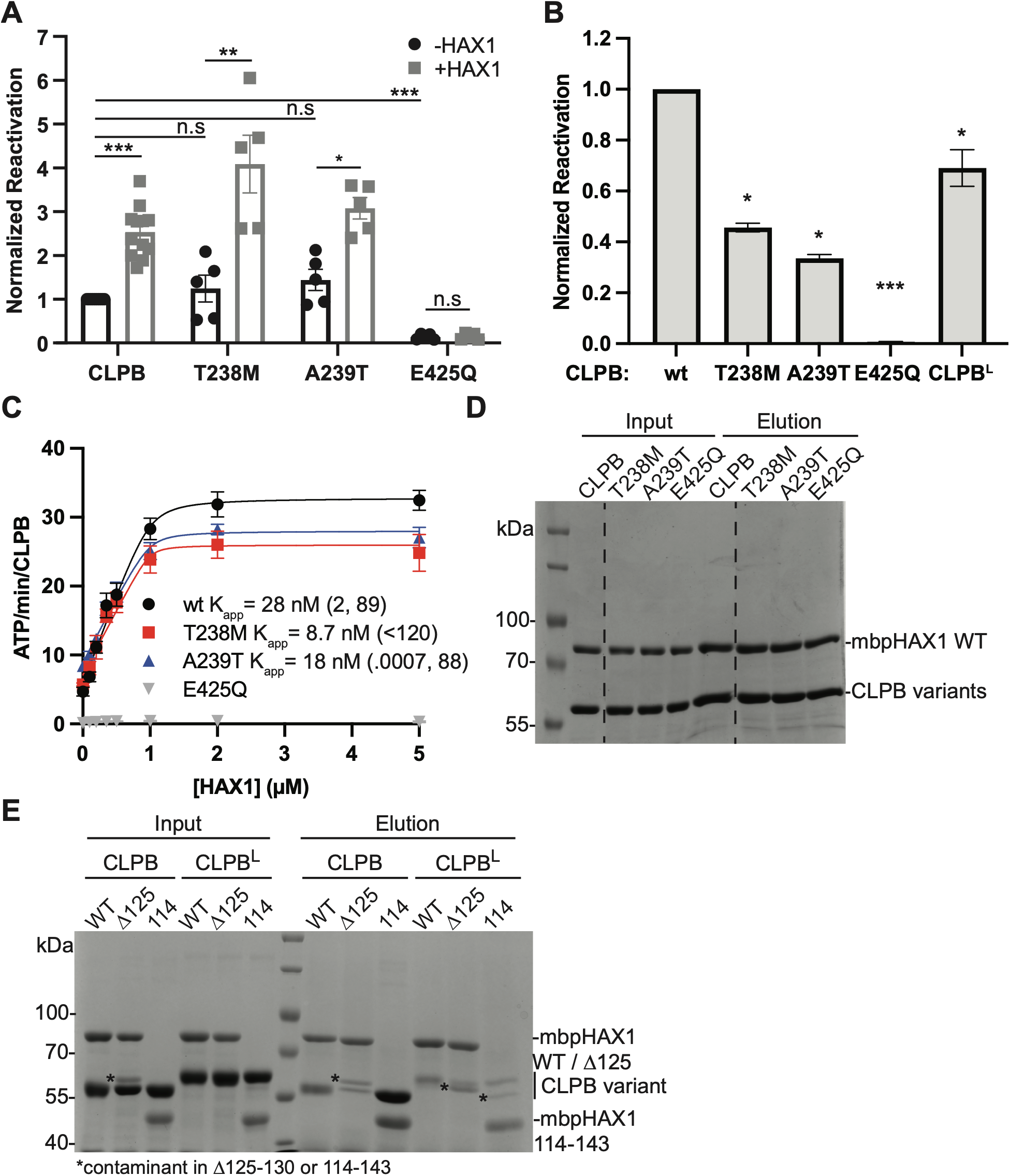
(A) Disaggregation of FFL_agg_ by 0.5 μM CLPB variants ± equimolar HAX1 after 90 min, n = 5 for CLPB variants and 10 for CLPB. (B) Disaggregation of FFL_agg_ by 5 μM CLPB variants after 90 min; n = 2 for CLPB and point variants; n = 3 for CLPB^L^. P values denote difference from wildtype CLPB. (C) ATP hydrolysis rate of 1 μM CLPB variants over HAX1 concentration (0-5 μM), n = 2 for variants and 12 for wildtype CLPB. (D, E) Pulldown results of HAX1 (bait) with different CLPB variants, n = 3. For all plots, error = SEM. Statistical significance denoted by p < 0.05 (*), p < 0.005 (**), p < 0.0005 (***).

**Figure S4, related to Figure 4.**
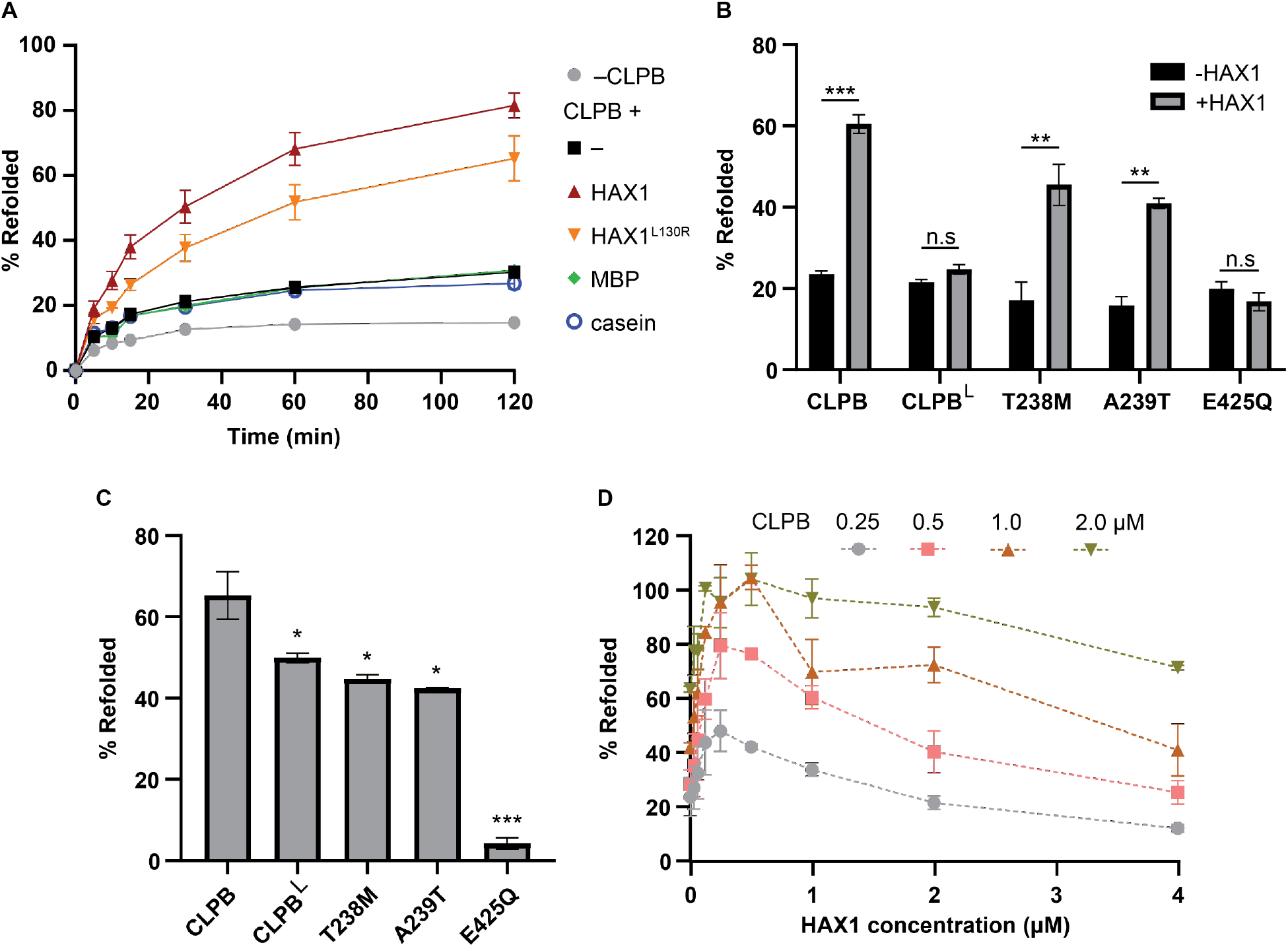
(A) The effect of 0.5 μM HAX1 on 0.5 μM CLPB refoldase activity compared to 0.5 μM HAX1 variants or protein controls, n ≥ 4. (B) The effect of 0.5 μM HAX1 on the refoldase activities of 0.5 μM CLPB variants, n ≥ 3. (C) Basal refoldase activity of CLPB variants at 5 μM, n = 2 for CLPB^T238M^ and CLPB^A239T^, 4 for CLPB^E425Q^, 6 for CLPB, and 8 for CLPB^L^. (D) HAX1 (0-4 μM) effect on the refoldase activity of CLPB over several concentrations (0.25-2.0 μM CLPB) after 90 min; n = 2. Error = SEM for all plots. Statistical significance denoted by p < 0.05 (*), p < 0.005 (**), p < 0.0005 (***).

**Figure S5, related to Figure 5.**
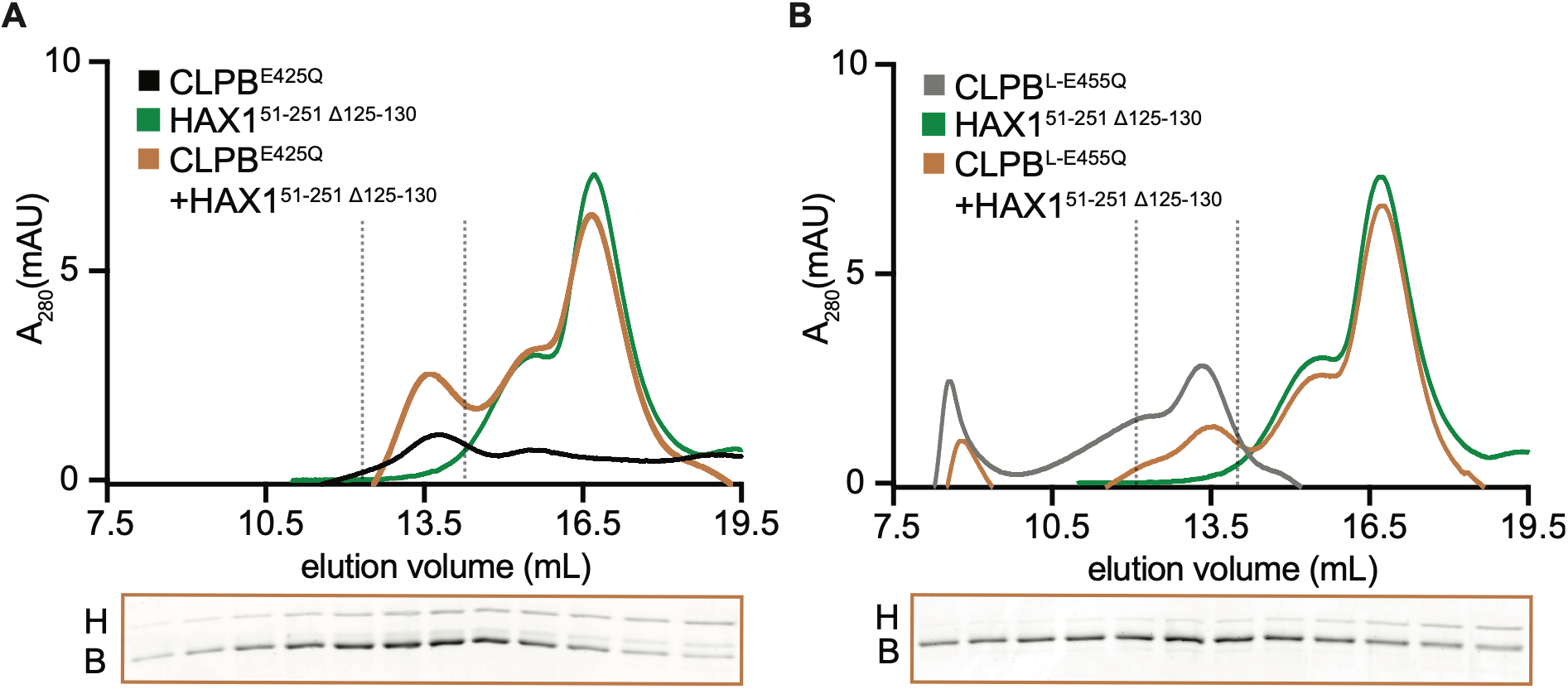
Size exclusion chromatography to assess CLPB, HAX1, and CLPB-HAX1 complex formation with (A) CLPB^E425Q^ and HAX1 51-251 Δ125-130; (B) CLPB^L-E455Q^ and HAX1 51-251 Δ125-130. Elution fractions of CLPB+HAX1 highlighted between the vertical dashed lines were analyzed by SDS-PAGE and Sypro Orange staining and are displayed below each chromatogram. For all chromatograms, n=1.

